# Machine learning-enabled cancer diagnostics with widefield polarimetric second-harmonic generation microscopy

**DOI:** 10.1101/2021.05.26.445874

**Authors:** Kamdin Mirsanaye, Leonardo Uribe Castaño, Yasmeen Kamaliddin, Ahmad Golaraei, Renaldas Augulis, Lukas Kontenis, Susan J. Done, Edvardas Žurauskas, Brian C. Wilson, Virginijus Barzda

## Abstract

The extracellular matrix (ECM) collagen undergoes major remodeling during tumorigenesis. However, alterations to the ECM are not widely considered in cancer diagnostics, due to mostly uniform appearance of collagen fibers in white light images of hematoxylin and eosin-stained tissue sections. Polarimetric second-harmonic generation (P-SHG) microscopy enables label-free visualization and ultrastructural investigation of non-centrosymmetric molecules, which, when combined with texture analysis, provides multiparameter characterization of tissue collagen. This paper demonstrates whole slide imaging of breast tissue microarrays using high-throughput widefield P-SHG microscopy. The resulting P-SHG parameters are used in classification to differentiate tumor tissue from normal with 94.2% accuracy and F1-score, and 6.3% false discovery rate. Subsequently, the trained classifier is employed to predict tumor tissue with 91.3% accuracy, 90.7% F1-score, and 13.8% false omission rate. As such, we show that widefield P-SHG microscopy reveals collagen ultrastructure over large tissue regions and can be utilized as a sensitive biomarker for cancer diagnostics and prognostics studies.

## Introduction

Cancer is amongst the leading causes of death, affecting approximately 1 in 5 people worldwide; a figure that is predicted to increase by ∼ 47% by 2040 [1]. The most common cancer diagnostic techniques rely on the gold standard histopathology of hematoxylin and eosin (H&E) stained tissue sections examined with white light microscopy. H&E histopathology focuses predominantly on characteristics of the cell nuclei and the tissue cell arrangements. However, the ultrastructure and texture of the background collagen-rich extracellular matrix (ECM) can also serve as an additional biomarker.

Collagen is a major constituent of the ECM, which undergoes structural alterations during tumorigenesis [2]. Several stains, including Movat’s pentachrome [3], Masson’s trichrome [4], picrosirius red [6], as well as immunohistochemical labels [5] have been used to highlight collagen. However, the use of ECM collagen as a biomarker is not widely employed in cancer diagnostics due to subtle structural variations which may be too difficult to detect with white light microscopy.

Second-harmonic generation (SHG) microscopy provides an alternative imaging modality, which enables label-free visualization of the collagenous ECM with high specificity [7, 8]. The SHG signal depends on the inherent 3D structure of noncentrosymmetric collagen fibrils, characterized by the nonlinear susceptibility tensor, as well as incident laser polarization [9]. By taking advantage of SHG intensity’s dependence on laser polarization, polarimetric SHG (P-SHG) can be utilized in laser scanning microscopy to measure the nonlinear susceptibility tensor elements for each imaged voxel of the tissue. It was further used to identify cancer-associated ECM alterations of multiple human tumor types in lung [10], thyroid [11], breast [12, 13, 14], pancreas [15], and ovary [16]. However, current P-SHG microscopy techniques rely on raster scanning of the imaged area, which are slow for whole slide imaging and high-throughput clinical use.

Recently, widefield SHG microscopy was shown to significantly reduced the time required for large-area imaging compared to laser scanning systems [17]. Based on this technology, we have developed a high-throughput quantitative imaging technique by integrating P-SHG with widefield microscopy. Widefield P-SHG microscopy enables rapid scan-less imaging of large sample regions (∼ several millimeters) using 16 orthogonal polarization states. Moreover, the subsequent image processing avoids time consuming pixel-by-pixel model fitting, and provides a series of polarimetric parameters that highlight the ultrastructural properties of the collagenous ECM. Furthermore, morphological organization of collagen fibers in the ECM of cancer tissue can be investigated using texture analysis of polarimetric parameters [18, 19]. Texture analysis of P-SHG polarimetric parameters was previously utilized in studies of lung cancer, revealing significant features that aid in characterization of tumor tissue [20].

In this work, polarimetric and texture parameters from widefield P-SHG imaging were used to train a logistic regression classifier, which was able to differentiate normal from tumor tissue with 94.2% accuracy and F1-score, and 6.3% false discovery rate, in human breast tissue microarray slides. The trained classifier was then further evaluated by predicting the presence of tumor on an independent data set, yielding 91.3% accuracy, 90.7% F1-score, and 13.8% false omission rate. Implementation of machine learning into the image postprocessing enhances differentiation of normal and tumor tissues, potentially enabling automated screening of, for example, tissue microarrays, as well as mapping areas of altered collagen to improve histopathologic diagnoses.

## Results

### Experimental design and overview

We have developed a widefield P-SHG microscopy technique for rapid high-throughput quantitative imaging, which we applied to breast tissue microarrays. The method generates a set of information-rich polarimetric and texture parameters, and utilizes these to classify normal and tumor tissues.

Figure 1 shows an overview of the experimental workflow. The tissue microarray slide was imaged under widefield P-SHG microscopy with 16 unique combinations of input and output polarization states (Fig. 1a). These were combined to generate images of SHG Stokes vector elements, in accordance with reduced double Stokes-Mueller polarimetry, as illustrated in Fig. 1b (see supplementary information for more details). Images of the following 5 distinct polarimetric parameters were extracted and used to characterize the ultrastructure of collagen in each image pixel: 1) average SHG intensity from all incident circular polarizations, 2) R-ratio, which is the ratio of 2 achiral second-order nonlinear optical susceptibility elements (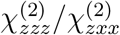, where the z-axis is parallel to the collagen fiber axis), 3) degree of circular polarization (DCP), 4) SHG circular dichroism (SHG-CD), and 5) SHG linear dichroism (SHG-LD). Each polarimetric parameter carries unique ultrastructural information about the ECM collagen.

**Figure 1:**
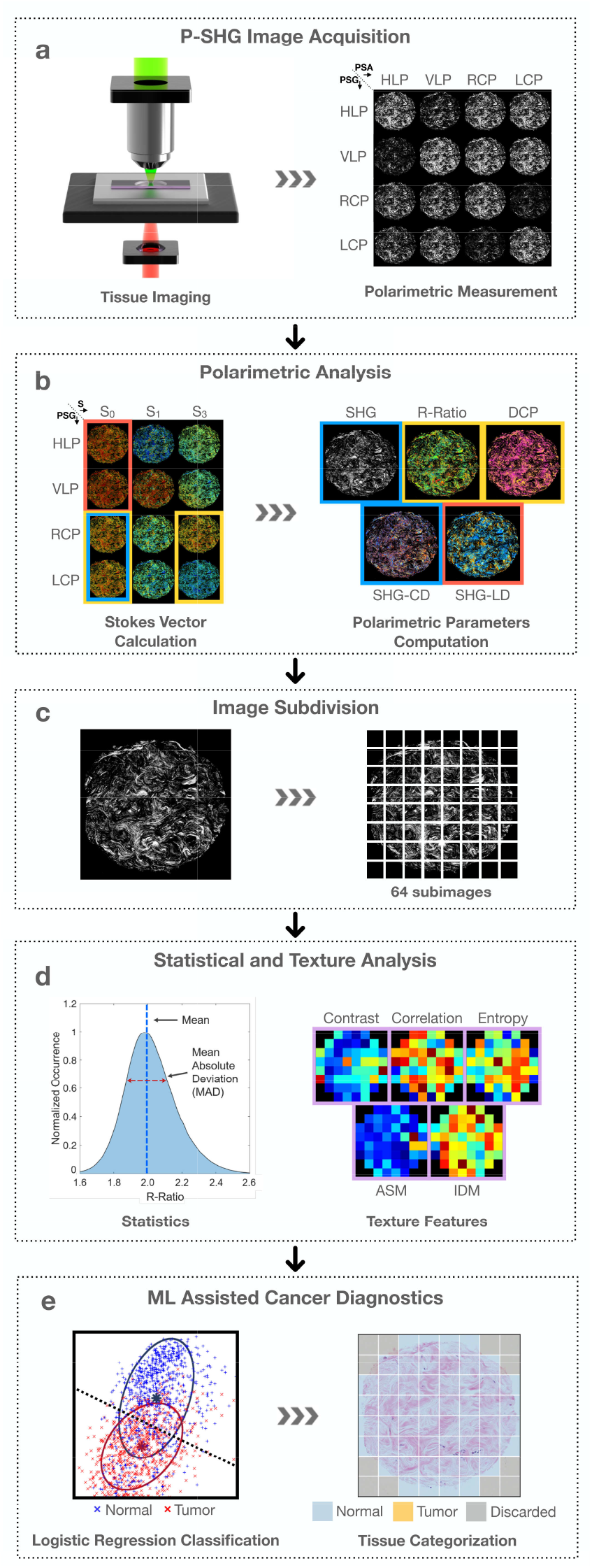
From imaging to classification. a, Widefield polarimetric SHG imaging of the sample at 16 unique polarization state combinations. b, Calculation of SHG Stokes vector elements to compute polarimetric parameter images. c, Subdivision of polarimetric images into 64 sub-images, to allow high-resolution texture analysis and statistical significance testing. d, Calculation of mean and mean absolute deviation of polarimetric parameter, as well as contrast, correlation, entropy, angular second moment (ASM), and inverse difference moment (IDM) texture parameters of each sub-image e, Training of a logistic regression classifier using the polarimetric and texture parameters, and the subsequent prediction to differentiate normal and tumor tissue.

The SHG intensity obtained with circular incident polarization is highly sensitive to the molecular organization of collagen and is independent of the in-plane orientation of collagen fibers. It has been shown that the SHG intensity is reduced in various solid tumors [21, 22]. The R-ratio describes the structural organization of the collagen fibers, and has been successfully used to differentiate normal and malignant tissues in lung, breast, thyroid, and pancreas [10, 12, 11, 15]. The value of DCP is closely related to the R-ratio in non-scattering tissue and reflects the disorder and depolarization in the sample. SHG-CD provides information on the collagen polarity and the out-of-plane fiber orientation in the tissue [23, 24, 9], which has been previously utilized in investigations of ovarian cancer [25]. Here, we introduce SHG-LD as a new parameter in P-SHG microscopy to study the in-plane organization of collagen fibers in the ECM. A similar definition of SHG-LD was previously used in surface SHG measurements of Langmuir–Blodgett films of chiral polymers functionalized with a nonlinear optical chromophore [26]. These five parameters provide complementary information on the ultrastructure of tissue collagen.

The images of each computed polarimetric parameter were further subjected to texture and statistical analyses. Texture analysis is a well-known method for characterizing tissue morphology by analyzing the variation in the neighboring pixel values [27]. It is commonly used as a classification tool through its ability to recognize patterns that may be indistinguishable to the human eye [28, 29, 30]. Here, each polarimetric image is comprised of over 4 million pixels. Hence, to enable high-resolution mapping of texture parameters and statistical investigations, all polarimetric images were divided into 64 sub-images (Fig. 1c). The mean, mean absolute deviation (MAD), and 5 of the most useful textures of the polarimetric parameters, including contrast, correlation, entropy, angular second moment (ASM), and inverse difference moment (IDM) were computed over all sub-images to characterize the collagen in the tissue (Fig. 1d) [18, 19].

The final stage of the analysis involved machine learning-assisted diagnostics, as shown in Fig. 1e. For this, the combination of mean, MAD and texture of 5 polarimetric parameters of normal and tumor tissue were used to train a logistic regression classifier. The trained classifier was used to perform predictions on an independent set of images and map the ultrastructural properties of collagen, leading to differentiation of normal and tumor tissue.

### Whole slide P-SHG imaging of tissue microarray

The collagen content of a breast tissue microarray was imaged using widefield P-SHG microscopy, resulting in images of whole breast tissue cores that were 0.6 mm in diameter, without the need for raster scanning. Figure 2a shows the annotated H&E-stained microarray of normal (N) and tumor (T) cores. The latter were characterized by estrogen receptor (ER), progesterone receptor (PR) and human epidermal growth factor receptor 2 (HER2) expression, and 3 of the most common subtypes were considered: ER+/PR+/HER2+ (triple positive or +/ + /+), ER+/PR+/HER2− (double positive or +/ + /−), and ER− /PR− /HER2− (triple negative or − / − /−) [31].

**Figure 2:**
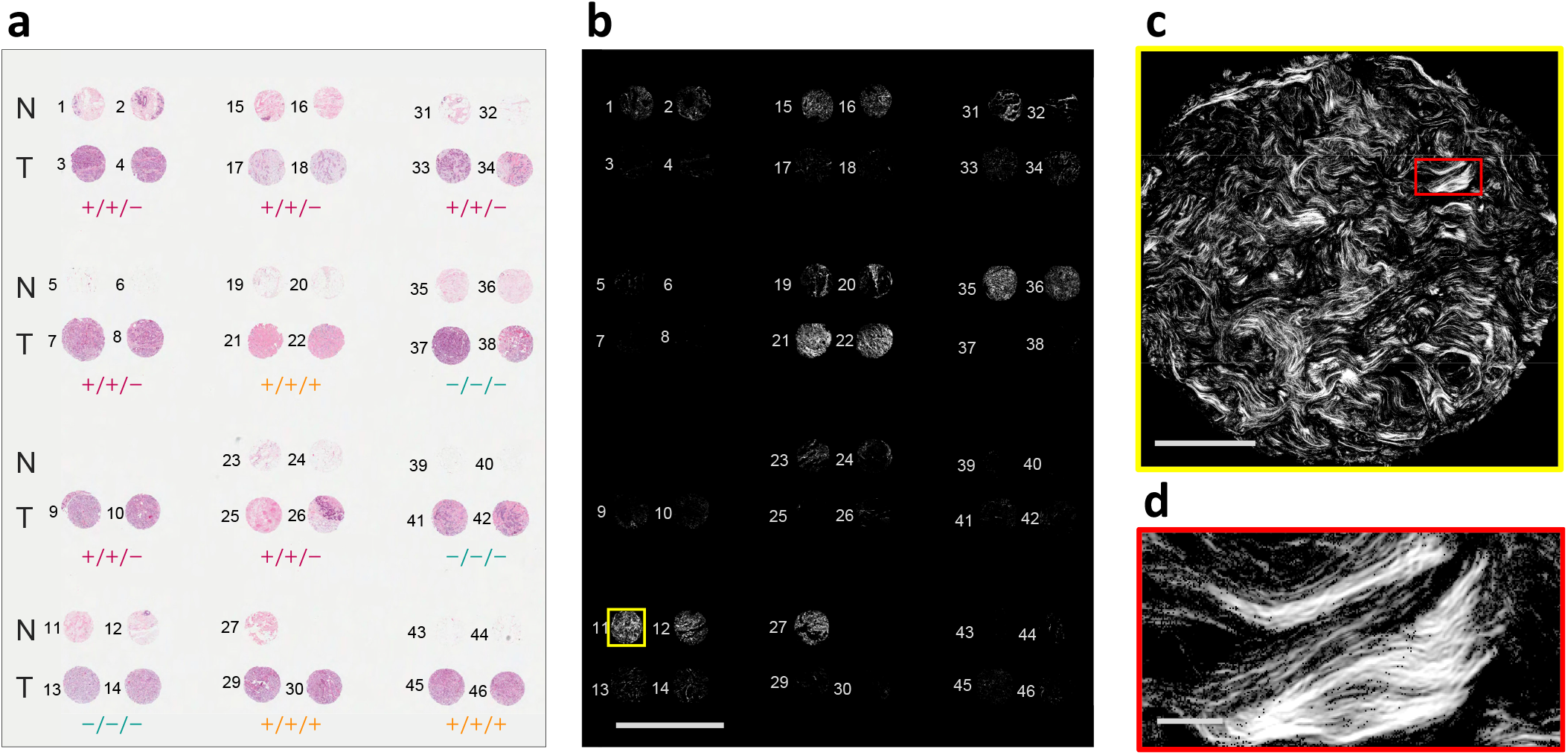
High-throughput widefield P-SHG imaging of tissue microarray. a, A H&E-stained breast tissue microarray containing normal and three distinct tumor subtypes. The positive sign indicates overexpression of estrogen receptor (ER), progesterone receptor (PR), and human epidermal growth factor receptor 2 (HER2), denoted by ER/PR/HER2. b, Image of circularly-polarized SHG intensity of the entire microarray slide shows distinctive low signal associated with tumor cores, except cores 21 and 22 that possess a large amount of collagenous stroma. Scale bar: 2.5mm. c, A single widefield SHG image consisting of 2048×2048 pixels and an area of approximately 670*µ*m×670*µ*m. Scale bar: 200*µ*m. d, A magnified region of the tissue, highlighted by red rectangle in c shows high resolution of the imaging technique. Scale bar: 25*µ*m.

The SHG intensity images of all core are seen in Fig. 2b. Each core image is comprised of 2048×2048 pixels, corresponding to an area of approximately 670*µ*m×670*µ*m, as shown in Fig. 2c, which corresponds to core 11, as indicated by yellow border in Fig. 2b. The resolving power of the microscope (approximate pixel size of 0.3*µ*m×0.3*µ*m) is shown by an enlarged rectangular region in Fig. 2d, whose location is highlighted with the red rectangle in Fig. 2c. For better viewing of the SHG images, pixels that exhibited signal-to-noise ratio (SNR) <1 were removed. It is clearly seen that tumor tissue has markedly lower SHG intensity than normal, consistent with previous reports in multiple tumor types [10, 12, 11, 15, 32]. This is likely due to collagen degradation in the tumor microenvironment which results in randomly oriented fibrils, so resembling a centrosymmetric arrangement. It is also clear that there are considerably fewer pixels with SHG signal present in the tumor tissues, rendering pixel count as an important classification parameter.

### Polarimetric SHG imaging and analysis

Representative polarimetric images of normal, triple negative, double positive, and triple positive groups are shown in Fig. 3. The H&E-stained core images (Fig. 3a) were segmented to reveal stained fibrillar components of the ECM using a published convolutional neural network (CNN) based technique [33] (Fig. 3b). H&E segmented images serve as a visual approximation of the total collagen content of the tissue. The true arrangement of the ordered collagen fibers can be visualized with P-SHG microscopy as presented in Fig. 3c. SHG identifies only the ordered collagen structures in the tissue; hence, highlighting a subset of the segmented collagen image. As shown in Fig. 3d, the tumor cores have larger R-ratio values than normal, similar to earlier reports of human breast tissues imaged with a scanning P-SHG microscope [12]. In addition to the extracted R-ratio, DCP of the SHG signal is obtained. As shown in Fig. 3e, the measured DCP was on average lower in normal tissue compared to each of the tumor groups.

**Figure 3:**
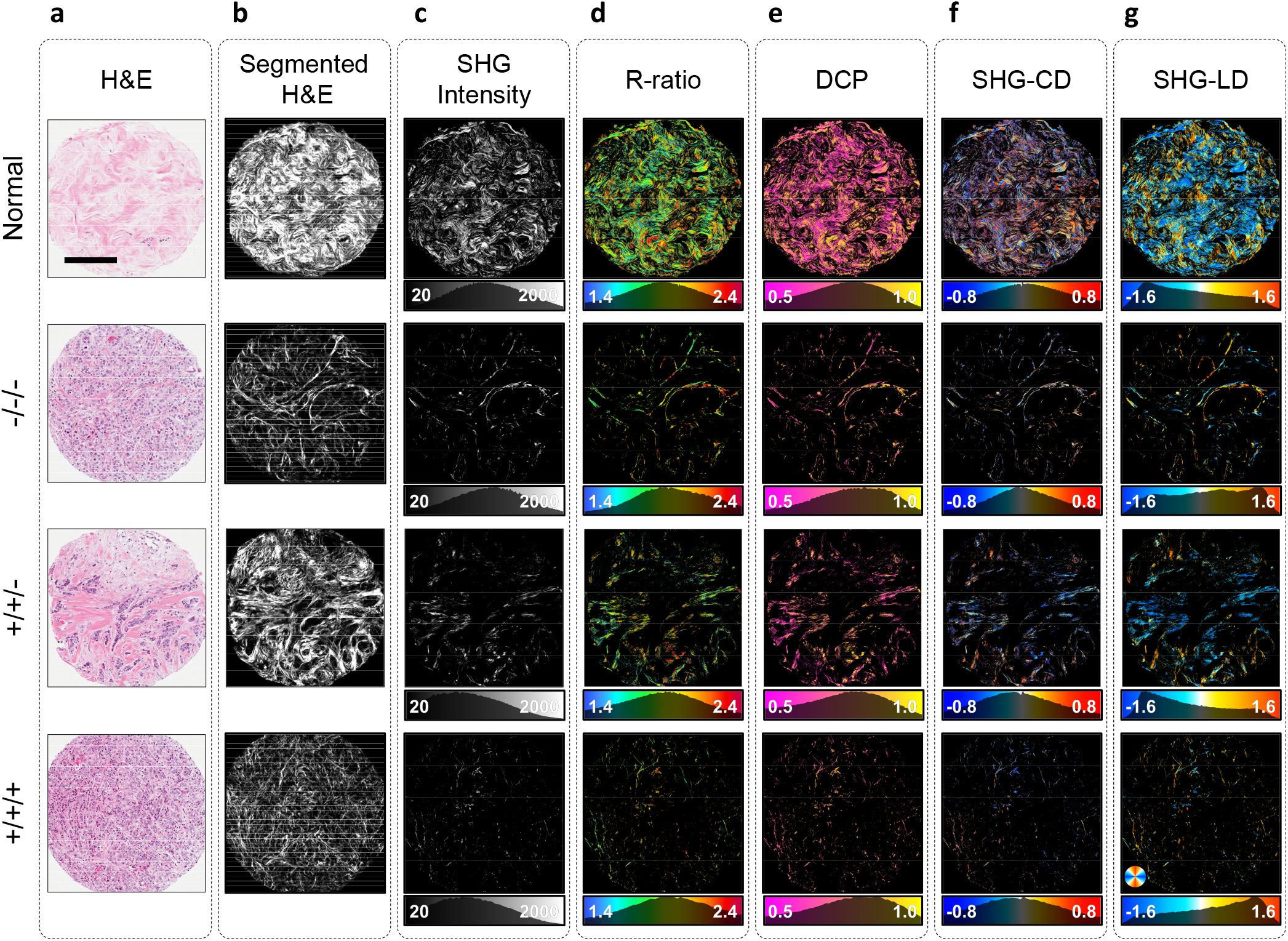
Widefield P-SHG polarimetric parameters. a, Representative H&E white light images of normal and three tumor breast cores with different expressions of the receptors ER/PR/HER2. Scale bar: 200*µ*m. b, Segmented images display the portion of H&E images stained by Eosin, which serves as a good estimate for total collagen content of the tissue. c, SHG intensity of circularly-polarized incident light, highlighting ordered collagen fibers. Normal core produces much larger SHG signal. d, R-ratio images of cores range from 1.4 to 2.4, indicating the presence of collagen in the tissue. e, Degree of circular polarization images show larger means in tumor due to higher R-ratio and low depolarization. f-g, SHG-CD and SHG-LD images depict the out-of-plane and in-plane fiber orientations, respectively. Due to unbiased sectioning of the tissue, collagen fibers are at random orientations, resulting in SHG-CD and SHG-LD distributions centered around zero.

SHG-CD values (Fig. 3f) were calculated using polarimetric images with circular input polarizations, and are directly associated with chiral and achiral optical susceptibilities, as well as the out-of-image-plane tilt angle of collagen fibers [9, 24]. Tissue samples were sectioned without any fiber orientation bias, resulting in random direction of collagen fibers. Therefore, SHG-CD mean values did not provide useful information for classification of tumor tissue. However, collagen in normal tissue appears wavier, leading to a broader distribution of the SHG-CD. Thus, MAD of SHG-CD distribution was instead used to differentiate between normal and tumor tissues. The SHG-LD images were constructed using the linear input polarizations (Figs. 3g), and highlight the in-plane collagen fiber orientation. Similar to SHG-CD, the SHG-LD distribution appears wider in normal tissue compared to the tumor, reflecting larger variations of the in-plane collagen fiber orientation in normal tissue. The SHG-LD measurements exhibited diverse distribution forms, due to random sectioning of breast cores. Hence, in order to further analyze the degree of variability of in-plane collagen fiber orientation, the MAD of SHG-LD distributions were computed and compared between normal and tumor groups.

### Texture analysis and statistical testing

Statistical multiple comparison tests were performed to evaluate the significance of parameter value differences between tumor and normal cores. Each core image was comprised of over 4 million pixels, however, pixels with SNR < 3 were discarded from analysis to ensure reliability of the statistical testing. Each large-area image of the cores was subdivided into 64 sub-images, each with 256×256 pixels (approximately 84*µ*m×84*µ*m in area). Sub-images located in the corner of each breast core P-SHG image, as well as those without viable signal were removed from the analysis. The number of pixels with SNR *>* 3 in each sub-image was used as an important metric, referred to as the SHG pixel density (PD) to indicate the abundance of ordered SHG-producing collagen in the tissue. The PD was found to be on average considerably higher in normal tissue than tumor; therefore, it was included in the set of diagnostic polarimetric and texture parameters. The mean, MAD, and texture features of the polarimetric parameters were used in multiple comparisons testing and machine learning-assisted classification. Given the large number of resulting parameters (36), the significance test results are shown as a circular plot in Fig. 4a. The circular plot is divided into four quadrants, representing normal, triple negative, double positive, and triple positive groups. The mean, MAD, and textures of each polarimetric parameter are highlighted by distinct colors. Most differences between normal and tumor groups were statistically highly significant (*p<* 0.001), as indicated by solid colored links. Links between tumor groups are mostly dashed lines, which indicate significant differences (0.001 *< p <* 0.05). Non-significant differences are omitted from the diagram. It is evident that all three considered molecular subtypes of breast cancer are well-differentiated from normal with comparable levels of accuracy using widefield P-SHG.

**Figure 4:**
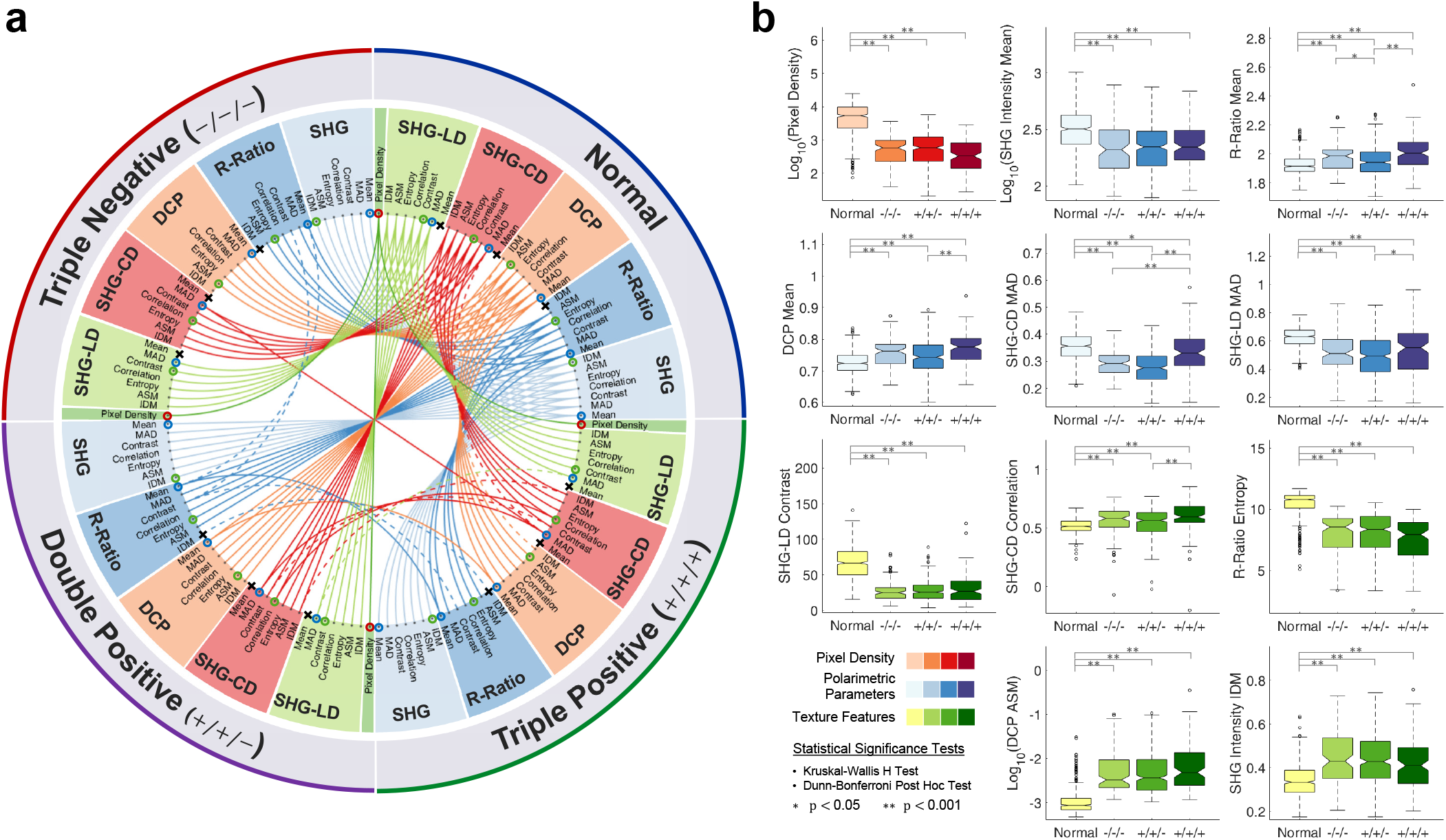
Parameter multiple comparisons testing. a, Circular plot of all multiple comparisons test between all four groups. Kruskal-Wallis H test and Dunn-Bonferroni post hoc test were performed. The dashed links indicate significant differences (0.001 *< p <* 0.05), while solid links depict highly significant differences (*p <* 0.001). All non-significant differences are omitted. It is evident that normal tissue is highly significantly different from all tumor groups across most of the measured parameters. Parameters that are not significantly different between normal and any of the tumor groups are denoted by x’s and removed from further investigation. b, Boxplots of selected parameters show detailed differences between all four considered groups. Color of the plots correspond to small circles around parameters in a. It is evident that triple positive is the most different tumor group from normal.

For a more detailed presentation of the data, boxplots of subsets of the computed parameters are shown in Fig. 4b. Sub-image means of both R-ratio and DCP showed similar trends of being significantly higher in tumor than normal tissue, with triple positive breast cancer possessing the largest R-ratio and DCP. The R-ratio results are in agreement with a previous investigation on small breast tissue regions [12]. The MAD of both SHG-CD and SHG-LD highlighted larger variation of the collagen fibers orientation in normal tissue, compared to tumor tissue. Between tumor groups, triple positive had the largest variation in fiber orientation, approaching that of the normal group. Three of the parameters (SHG-CD mean, SHG-LD mean, and R-ratio IDM) were not significantly different between normal and any of the tumor groups, as indicated by **x**’s on their corresponding nodes in Fig 4a. As such, these parameters were omitted from classification.

### Classification and prediction of normal and tumor breast tissue

To showcase the efficacy of collagen as a diagnostic biomarker and display the detection capabilities of widefield P-SHG microscopy, the polarimetric parameter statistics (mean and MAD) and texture features were used to train a binary logistic regression classifier. Prior to classifier training, two cores (one from normal group and one from tumor group) were removed to form a prediction dataset, which was used to further investigate the predictive power of the classifier. All tumor cores were combined into a single tumor group and by treating each sub-image as an individual data point, 544 normal and 543 tumor data points (samples), across 33 different polarimetric and texture parameters (predictors) were used for classification. The training was repeated 1000 times with random stratified partitioning of the dataset for 5-fold cross-validation, and the standard deviation of performance metrics were used to evaluate the classifier stability [34].

Using the complete dataset, the classifier differentiated tumor from normal tissue with 94.2% accuracy at 50% posterior probability threshold, as indicated in Fig 5a. The threshold-dependent classification performance metrics (true negative rate, true positive rate, negative predictive value, and positive predictive value) were within 93-95%. The complimentary metrics (false positive rate, false negative rate, false omission rate, and false discovery rate) were within 5-7%. In addition, the F1-score of 94.2% demonstrates the excellent robustness and accuracy of the trained classifier in identifying tumor tissue regions.

**Figure 5:**
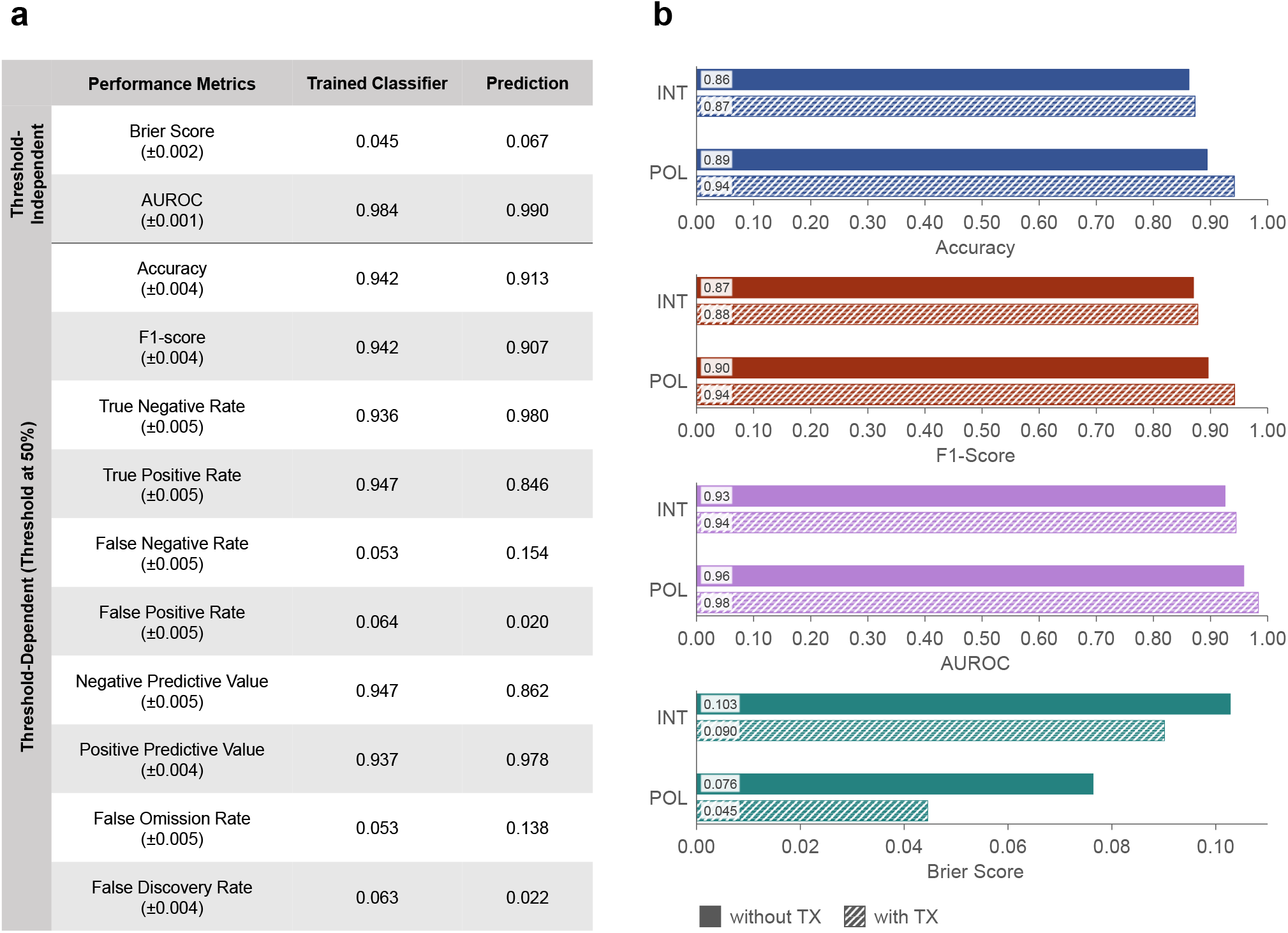
Tissue classification performance. a, Tissue classification and prediction with a binary logistic regression classifier, trained using all polarimetric and texture parameters. Reported values are mean ± standard deviation. b, Testing the impact of various parameters using 4 subsets of the measured parameters, including SHG intensity (INT) and all 5 polarimetric parameters (POL), with and without texture parameters (TX). All groups include pixel density. Differences in accuracy, F1-Score, AUROC, and Brier score between groups are statistically highly significant (*p* 0.001), as determined by the Kruskal-Wallis H test and the Dunn-Bonferroni post hoc test. Using all polarimetric and texture parameters improves the classification performance across all metrics. All reported accuracy and F1-score values have < 0.4% standard error, and AUROC and Brier score have standard errors of < 0.2% and < 4.5%, respectively.

Two threshold-independent metrics, namely the area under the curve of the receiver operating characteristic (AUROC) and Brier score, were computed to further examine the classifier performance. An AUROC value of 1 is ideal, while 0.5 indicates a completely random classification. The Brier score ranges from 0 (perfectly accurate) to 1 (completely inaccurate). As shown in Fig 5a, the trained classifier generated an AUROC of 0.984 and Brier score of 0.045, indicating well-differentiated normal and tumor groups and low misclassification rate.

To determine the importance of polarimetric and texture parameters in classifier performance, 4 subsets of the data were considered (Fig 5b): 1) SHG intensity and PD, which are the most commonly used parameters, 2) SHG intensity and PD, together with corresponding texture analysis, 3) the 5 polarimetric parameters (including SHG intensity) and PD, and 4) the 5 polarimetric parameters and PD with corresponding texture parameters (complete dataset). Classification using only the SHG intensity resulted in the lowest accuracy, F1-score and AUROC values, and the largest Brier Score. Adding texture analysis provided a modest improvement of (∼ 1%) across all metrics. A more significant improvement was evident from using all 5 polarimetric parameters (including the SHG intensity), with accuracy and F1-scores reaching *>* 90%. Finally, combining all polarimetric and texture parameters, produced the best overall performance with accuracy and F1-score *>* 94%, AUROC *>* 98%, and Brier score of 0.045.

The trained classifier was then used on the prediction set of normal and tumor cores that had not been part of the training set. As depicted in Fig 5a, the prediction highlighted sub-images with normal and tumor properties, resulting in 91.3% accuracy, 0.990 AUROC, and 0.067 Brier score. Normal tissue was better predicted than tumor tissue, with a true negative rate of 98.0% compared to a true positive rate of 84.6%. However, the positive predictive value was 97.8%, resulting in an F1-score of 90.7%.

## Discussion

For the first time, we have presented the application of widefield P-SHG microscopy of the collagenous extracellular matrix as a potential histopathological biomarker for cancer diagnostics. The instrument provides rapid high-resolution large-area imaging of tissue samples such as the microarrays shown here, while providing comprehensive ultrastructural information through 33 quantifiable parameters. The lack of moving optics, such as scanning mirrors, in widefield imaging significantly contributes to the robustness of the technology. This enables simple construction and customization of the microscope, that is well suited for user-friendly high-throughput applications. Overall trends in many of the measured parameters showed a clear distinction between tumor and normal tissues. In particular, triple positive tissue had the greatest differences from normal tissue across most parameters. However, this distinction was small compared to the overall difference between normal and all tumor groups, as further illustrated by multiple comparisons tests on the circular plot of Fig. 4a.

Classification enabled an efficient use of all measured parameters, allowing for accurate predictions of normal and tumor tissue with performance comparable to other methods involving machine learning-assisted cancer diagnostics [35, 36, 37]. The pixel density and average SHG intensity were amongst the most informative measured parameters, with substantial differences in the ECM between normal and tumor tissues. It is important to note that simply the absence of SHG signal and, thus, lower pixel density and average SHG intensity are not always indicative of the presence of tumor; sparsely distributed collagen fibers and adipose tissue often do not result in sufficient SHG signal, therefore, such regions must be discarded to avoid inflation of false positive rates. Here, empty regions in the corners of the breast core images, and cores which were mainly comprised of adipose tissue were manually discarded. However, in future developments, H&E images may be used simultaneously with widefield P-SHG to identify and discard such areas automatically. Moreover, most breast cores showed a significant reduction in ordered collagen, as indicated by lower SHG intensity compared to normal tissue. However, 2 cores showed a high degree of stroma, and due to rarity of such tissue types in the microarray, they were discarded from classification training data.

The calculated classification metrics showed robust and accurate tissue differentiation with 94.2% accuracy, 6.3% false discovery rate, AUROC of 0.98, and Brier score of 0.045. Capabilities of the trained classifier were further demonstrated in predicting an independent test dataset with an accuracy of 91.3%, false omission rate of 13.8%, AUROC of 0.990, and a Brier score of 0.067. It is important to note that the number and size of the sub-images used, affect the predictive power of the classifier. For example, subdividing the images into fewer but larger sub-images would improve the accuracy and specificity, at the cost of decreased classification stability, resulting in reduced reliability. We found that subdivision level of 64 sub-images per image delivers highly accuracy classification accuracy with sufficient robustness. We present a detailed investigation of the optimum number of sub-images for classification of the P-SHG data in the supplementary information document (See Supplementary Fig. 2).

A natural extension to this work is introduction of chiral nonlinear susceptibility components. Consequently, additional polarimetric parameters such as chiral second order nonlinear optical susceptibility ratio 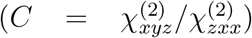) may be introduced to further improve molecular identification, based on varied polarity of the collagen fibers throughout the tissue sample. The C-ratio has been previously investigated in scanning P-SHG systems, revealing information on collagen fiber polarity and organization [9, 41].

In this study, we demonstrated widefield P-SHG microscopy as a potential tool in cancer diagnostics. However, it is important to also identify the applications of the microscope in studying other pathologies related to remodeling of the collagen structure in the ECM and in other collagenous tissues. As an example, widefield P-SHG microscopy may be used to identify large-scale ultrastructural changes in pathologies relating to abnormalities in fibrillar collagen types I, II, and III such as, arterial aneurysms, chondrodysplasias, osteogenesis imperfecta, osteoporosis, osteoarthrosis, intervertebral disc disease, and Ehlers-Danlos syndrome [38]. Scanning P-SHG has also been used to study muscle ultrastructure in rat and drosophila larvae [39, 40]. Future applications of the widefield P-SHG may include high-throughput investigations of human muscle pathologies such as muscular dystrophy and multiple sclerosis. In addition, continuous and uniform illumination of the entire imaged area provided by the widefield P-SHG microscope, enables dynamic imaging of processes involving fast kinetics, such as live ultrastructural imaging of contracting muscle fibers [17] and deformation of collagen fibers during application of external forces [44].

## Methods

### Widefield P-SHG microscopy

A custom microscope was designed and constructed for widefield polarimetric SHG imaging as shown in Fig 6. A high-power amplified laser (PH1-15W, Light Conversion) was used for large area illumination. The laser power was set to 1.3 W, at 100 kHz repetition rate, and 13*µ*J pulse energy. The laser beam was collimated to 4 mm diameter and coupled to the microscope. The beam passed through an infrared polarizing cube (PBS102, Thorlabs) to produce a linearly polarized state, followed by an infrared liquid crystal variable retarder (LCC1223-B, Thorlabs), referred to as the polarization state generator (PSG-LCVR), which was positioned with its fast axis at 45° to the incident linear polarization. An achromatic doublet (AC254-030-AB, Thorlabs) with a focal length of 30 mm was used to focus the beam on the sample. Widefield imaging was achieved by placing the sample above the focal plane and adjusting the illumination area (∼ 1 mm in diameter) through axial translation of the excitation achromatic doublet. A Plan-Neofluar 20×/0.50 air collection objective (420350-9900-000, Zeiss) was used to collect the SHG signal radiated in the forward direction. The polarization state analyzer (PSA-LCVR), comprised of a LCVR in the visible wavelength range (LCC1223-A, Thorlabs), was used to probe the outgoing polarization of the SHG signal (also positioned with its fast axis at 45° to the incident linear polarization). The SHG signal was passed through a tube lens (452960-0000-000, Zeiss) and a visible range polarizing beam splitter (PBS251, Thorlabs). A dichroic mirror (FF662-FDi02-t3-25×36, Semrock) was used to separate the visible SHG signal from infrared fundamental light. The infrared light transmitted through the dichroic mirror was blocked by a beam dump. The reflected SHG signal was filtered by two BG40 colored glass filters (FGB37-A, Thorlabs), a 515/10nm interference filter (65-153, Edmund Optics), and projected onto a sCMOS camera (ORCA-Flash 4.0 V2, Hamamatsu).

**Figure 6:**
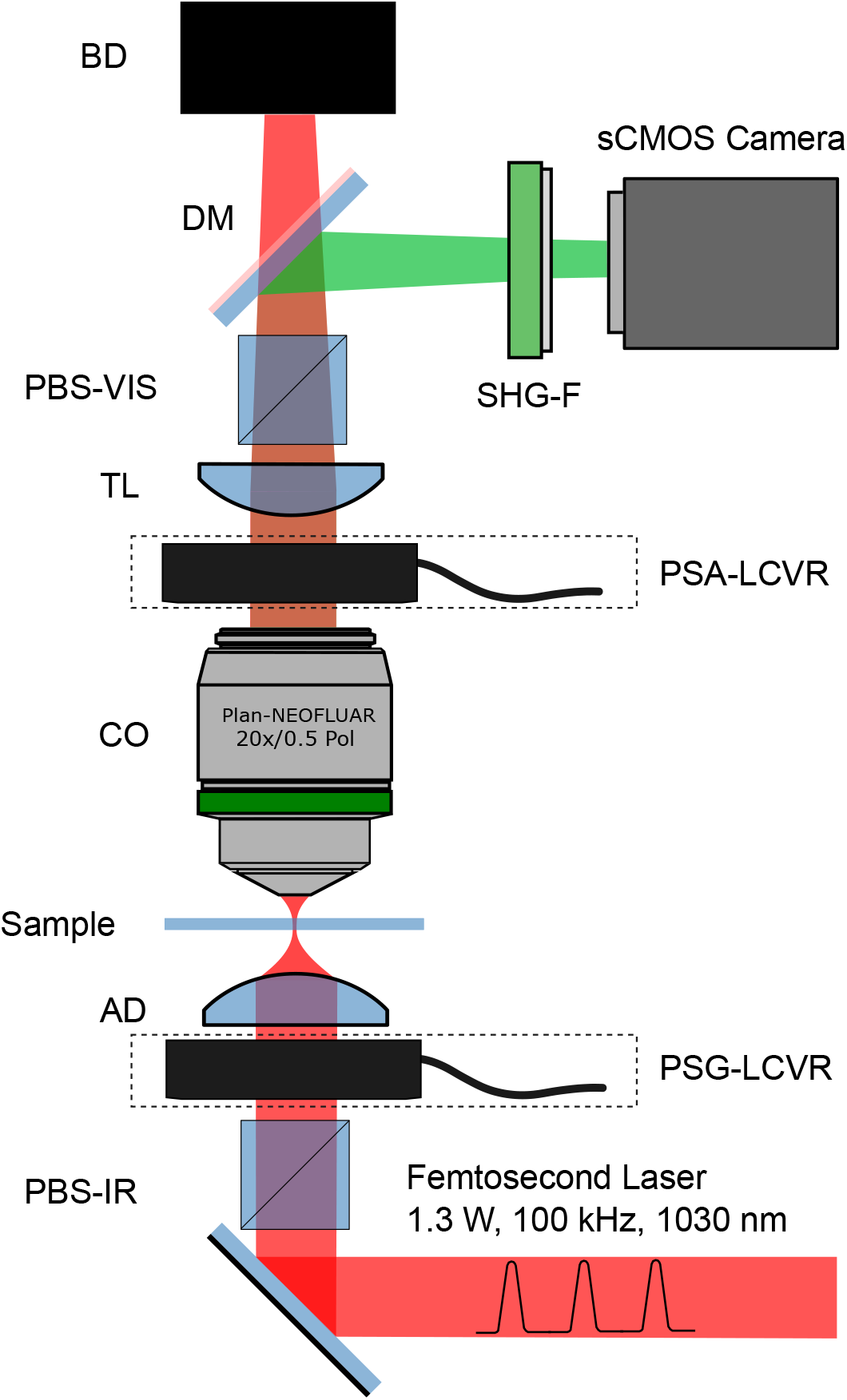
Widefield P-SHG microscope setup. Schematic representation of the experimental setup. The laser beam denoted in red is focused before the sample after passing polarizing beam splitter (PBS-IR), liquid crystal variable retarder of polarization state generator (PSG-LCVR) and achromatic doublet (AD). The produced SHG signal, depicted in green is collected by a collection objective (CO), passed through the liquid crystal variable retarder of polarization state analyzer (PSA-LCVR), tube lens (TL), polarization beam splitter (PBS-VIS), dichroic mirror (DM), filter (SHG-F), and projected onto the camera. The laser beam is terminated at a beam dump (BD).

To carry out polarimetric measurements, four orthogonal incoming polarization states were generated including left circular polarized (LCP), horizontally linearly polarized (HLP), right circularly-polarized (RCP), and vertically linearly polarized (VLP), corresponding to quarter-wave (λ/4), half-wave (λ/2), three-quarter-wave (3λ/4), and full-wave (λ) retardances of the PSG-LCVR, respectively. For each incoming polarization state, four SHG signal polarization states (set by the PSA-LCVR) corresponding to the same set of retardance values (quarter-wave, half-wave, three-quarter-wave and full-wave) were measured, resulting in 16 combinations of polarization states. Each polarization state was imaged with a 10 s exposure time, and 5 s of delay time was used for switching polarization states between exposure times, so that polarimetric measurement of each 670*µ*m×670*µ*m imaged area was completed in 4 minutes. Therefore, P-SHG imaging of the entire microarray was achieved in approximately 3 hours. It is important to mention that whole-microarray imaging time may be significantly reduced by decreasing the polarization switch delay time to <1s, in which case, whole-array P-SHG measurement would be complete in < 2.5 hours. Furthermore, the camera exposure time may be reduced with larger illumination areas and higher laser powers, however, such conditions were not considered in this work and will need to be tested for photobleaching and photodamage.

### Polarimetric parameter calculations

The polarization state of the SHG signal can be characterized by a Stokes vector [42, 43]

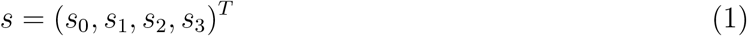

where *s*_0_ is the total intensity, *s*_1_, *s*_2_ are the linearly polarized Stokes vector components, and *s*_3_ is the circularly-polarized Stokes vector component. The measured SHG of the 16 polarization state combinations were used to compute SHG Stokes vector elements. Using these elements, polarimetric parameters of interest were calculated to reveal ultrastructural properties of interest, including: the average SHG intensity produced with circularly-polarized incident light; a ratio of two achiral second order nonlinear optical susceptibility tensor elements, 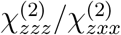 (R-ratio), where z points along the fiber axis; the degree of circular polarization (DCP); SHG circular dichroism (SHG-CD); and SHG linear dichroism (SHG-LD). R-ratio and DCP have been shown previously to provide valuable information on the ultrastructural and molecular organization of collagen fibers, while SHG-CD and SHG-LD probe the three-dimensional fiber orientation [10, 12, 11, 15, 41, 44, 45]. For detailed calculation of all the Stokes vector elements, refer to supplementary information document. The orientation-independent SHG, *I*^*CP*^, of each core was computed from the average SHG signal from right and left (RCP and LCP) circular polarizations of the incident light

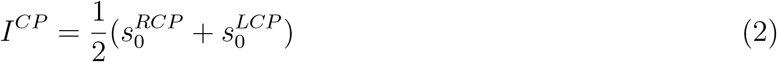

where the superscripts refer to the PSG polarization states, and the subscripts identify Stokes vector components of SHG signal. The R-ratio was calculated in terms of 1st and 4th elements of the measured SHG Stokes vector [48]:

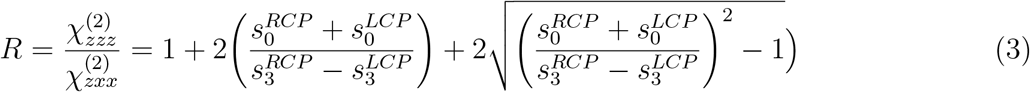

DCP was defined as the average of the magnitude of circular Stokes element, over the total intensity, such that [49]

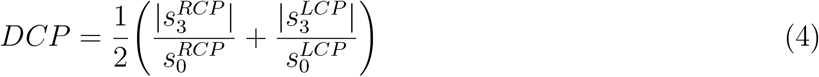

In addition, we computed SHG-CD, which was conventionally expressed as [46, 47, 48] and SHG-LD as:

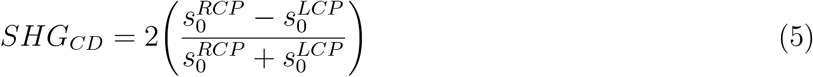

and SHG-LD as:

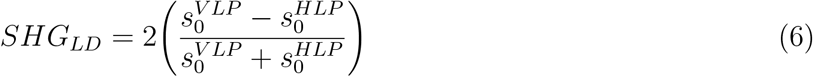

where both SHG-CD and SHG-LD range from -2 to 2.

### Texture analysis

Textural parameters were extracted from a grey-level co-occurrence matrix (GLCM), which is a second-order statistical representation of the grey-level distribution in a region of interest [27]. The GLCM was built by counting the occurrence of a pixel of grey level *i* followed by a pixel of grey level *j* at a distance *d* along a direction specified by angle *θ*. In the case of nearest neighbors (*d* = 1), the four angles of analysis were *θ* = {0°, 45°, 90°, 135°}. GLCMs of all four angles were constructed and averaged, forming a direction-independent GLCM. The probability density function, *P*_*d,θ*_(*i, j*), of finding certain grey-level pairs was obtained through normalization of the resulting GLCM. From this, we calculated the five textural parameters of interest, as described by Haralick et.al [27].

A custom-built texture analysis program was written in MATLAB, taking advantage of the available functions. The size of the GLCM depends on the number of grey levels (*N*_*g*_) specified by the user. Since the program runtime was highly dependent on *N*_*g*_, a testing routine was carried out to determine the optimal *N*_*g*_, represented by discretization level of continuous polarimetric parameters (refer to supplementary information and Supplementary Fig. 1 for detailed calculation of *N*_*g*_).

The presence of significant background in widefield P-SHG imaging results in highly skewed texture parameter distributions, leading to reduced differentiation between the groups. As such, the background pixels were converted to “not a number” (NaN) and were not included in the analysis. Consequently, the texture parameters only reflect the structure of the signal-producing entities, such as collagen fibers.

The measured texture parameters were contrast, correlation, entropy, angular second moment (ASM) and inverse difference moment (IDM). The contrast, *ν*, is defined by:

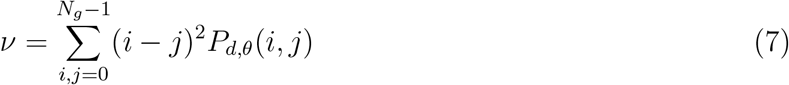

which is a measure of pixel value differences between grey-level pairs. Contrast is highly sensitive to variation in neighboring pixel values and it can be used to probe collagen fiber density in the tissue.

Correlation, *ρ*, quantifies a linear dependence between grey-level pairs, is expressed as:

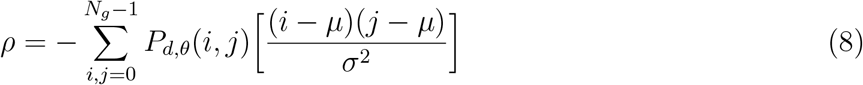

where *µ* is the mean and *σ* is the standard deviation of the grey levels. The correlation ranges from -1 to 1 for perfectly negatively or perfectly positively correlated images, respectively. In the context of widefield polarimetric SHG microscopy, correlation may be used to showcase well-defined structures and patterns in the acquired images. Entropy, *S*, is defined by:

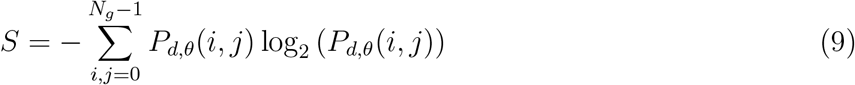

which measures the level of disorder or lack of spatial organization of the grey levels. Entropy increases as disorder increases; however, it is also indicative of the size of the pixel clusters, which often represent bundles of collagen fibers.

Conversely, angular second moment (ASM) is a measure of orderliness and ranges from 0 to 1, where 1 is characteristic of a completely uniform image. ASM also varies with the size of pixel clusters of comparable values and is a measure of the average size of similar collagen fiber bundles in an image. ASM is expressed as:

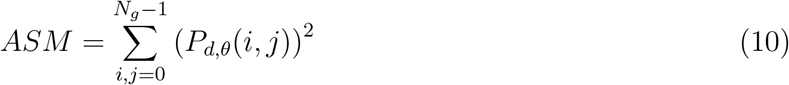

Finally, the inverse difference moment (IDM) describes the homogeneity of an image. Similar to ASM, the range of IDM is from 0 to 1, where 1 is indicates of a completely uniform image. IDM is provided by:

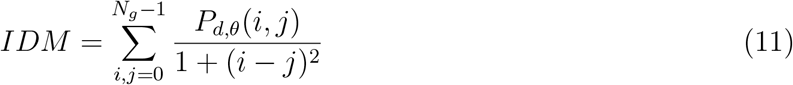

### Statistical analysis and machine learning

Each polarimetric image was divided into 64 sub-images for high-resolution texture analysis, and to compute the statistics of polarimetric parameters. The performance and stability of the logistic regression classifier in differentiating tumor from normal tissue were considered to establish the optimal discretization level of 64 sub-images for each polarimetric image (see supplementary information and Supplementary Fig. 2 for more detail). The results indicated a range of optimal discretization levels from 4 to 64 sub-images per image. The latter was chosen, since it results in a large AUROC and Brier scores, while performing with the highest stability.

Sub-images without SHG signal were discarded and the mean and mean absolute deviation (MAD) of all remaining sub-images were computed using standard MATLAB toolboxes. To reduce the effects of extreme outliers, data points below the 1st, and above the 99th percentiles of parameters in each group were discarded. These statistics were then used in further multiple comparisons tests to establish statistical significance between various groups. There were 544 normal, 103 triple negative, 325 double positive, and 115 triple positive data points across 36 polarimetric and texture parameters. Most parameters did not form normal distributions as indicated by Q-Q plots, and the Shapiro-Wilk test. Hence, MATLAB’s Kruskal-Wallis H test with 1086 degrees of freedom, along with the Dunn-Bonferroni post hoc test, were used to determine the significance of the difference between normal and all three tumor groups, for all polarimetric and texture parameters, separately. Differences with p-values < 0.05 and < 0.001 were considered to be significant and highly significant, respectively.

### Breast tissue microarray preparation

The tissue samples were collected following an institutionally-approved protocol (University Health Network, Toronto, Canada). The microarray contained 45 circular tissue samples (cores) from 12 different patients. The cores were 0.6 mm in diameter and 5 *µ*m thick, obtained from formalin-fixed tissues and mounted on a glass slide. The tissue was H&E-stained and imaged with a bright-field microscope scanner (Aperio Whole Slide Scanner, Leica Biosystems) for reference. Cores that were mainly comprised of adipose tissue did not produce significant SHG signal and were removed from the analysis. In addition, two tumor cores containing significant collagenous stroma with minor tumor foci were excluded, due to their rarity in the microarray. In total, 15 normal cores, 5 triple negative cores, 11 double negative cores, and 4 triple positive cores (20 tumor cores in total) were considered. The tissue assessment and tumor identification were performed by expert pathologists (S.J.D. and E.Z.).

## Acknowledgements

This work was supported by Natural Sciences and Engineering Research Council of Canada (NSERC) (RGPIN-2017-06923, DGDND-2017-00099, CHRPJ 462842-14), the Canadian Institutes of Health Research (CIHR) (CPG-134752), and European Regional Development Fund with the Research Council of Lithuania (01.2.2.-LMT-K-718-02-0016). We thank Light Conversion for providing the laser for experiments.

## Author Contributions

V.B. developed the concept of widefield P-SHG microscopy. L.K. optimized laser source for the setup. V.B., B.C.W., and K.M. designed the experiment. K.M., L.J.U., and V.B. constructed the widefield P-SHG microscope. S.J.D. prepared the breast tissue microarray. S.J.D. and E.Z. selected normal and tumor breast cores for P-SHG imaging. K.M. and L.J.U. calibrated the microscope and imaged the breast tissue microarray. K.M. developed the widefield P-SHG software, analyzed data, performed statistical significance testing, and carried out the classification. Y.K. developed the texture analysis software and analyzed data. R.A. performed image segmentation of H&E-stained tissue images. K.M., L.J.U., Y.K., A.G., and V.B. interpreted the results and wrote the article. All authors contributed to the manuscript.

