## Supplemental Information for "Machine learning-enabled cancer diagnostics with widefield polarimetric second-harmonic generation microscopy"

### Supplementary Note: Calculation of SHG polarimetric parameters

The polarimetric measurements were achieved using a polarization state analyzer (PSA) and a polarization state generator (PSG), according to double Stokes Mueller polarimetry (DSMP) [1]. Here, we introduce reduced DSMP, that enables fast computation of SHG polarimetric parameters without pixel-to-pixel model fitting.

Using reduced DSMP, the SHG polarimetric parameters were obtained by measuring four outgoing polarization states of the SHG signal with PSA, for each of the four incoming polarization states generated by PSG, resulting in 16 polarization state combinations of the PSA and PSG. The Stokes vector formalism is used on the SHG signal for calculation of the polarimetric parameters.

The SHG stokes vector is defined as:

$$s = (s_0, s_1, s_2, s_3)^T \tag{1}$$

, where $s_0$, $s_1$, $s_2$, and $s_3$ represent the total intensity, 0° and 90° linearly polarized, ±45° linearly polarized, and circularly-polarized components, respectively.

Polarization of the SHG signal measured by the camera, $s'$, can be describe by [2]:

$$s' = M_{PSA} s \tag{2}$$

, where $M_{PSA}$ represents the Muller matrix of the PSA, and $s$ is the polarization state of the generated SHG signal at the sample. The PSA contains a liquid crystal variable retarder (LCVR) and a polarizing beam splitter (PBS). The LCVR was placed at 45° with respect to the laboratory axis, resulting in the following Mueller matrix [3]:

$$M_{LCVR}(\delta) = \begin{pmatrix} 1 & 0 & 0 & 0 \\ 0 & \cos(\delta) & 0 & -\sin(\delta) \\ 0 & 0 & 1 & 0 \\ 0 & \sin(\delta) & 0 & \cos(\delta) \end{pmatrix} \tag{3}$$

, where $\delta$ represents the retardance. In addition, the PBS Mueller matrix in horizontal transmission configuration is provided by

$$M_{PBS} = \frac{1}{2}\begin{pmatrix} 1 & 1 & 0 & 0 \\ 1 & 1 & 0 & 0 \\ 0 & 0 & 0 & 0 \\ 0 & 0 & 0 & 0 \end{pmatrix} \tag{4}$$

Thus, the Mueller matrix of PSA ($M_{PSA}$) containing the LCVR and PBS is calculated as:

$$M_{PSA}(\delta) = M_{PBS}M_{LCVR}(\delta) = \frac{1}{2}\begin{pmatrix} 1 & \cos(\delta) & 0 & -\sin(\delta) \\ 1 & \cos(\delta) & 0 & -\sin(\delta) \\ 0 & 0 & 0 & 0 \\ 0 & 0 & 0 & 0 \end{pmatrix} \tag{5}$$

By treating $s$ as an input Stokes vector for the PSA, the outgoing Stokes vector, $s'$, is calculated using (2). Only the first component of $s'$ vector, corresponding to the full intensity of the SHG signal, is detected by the camera; hence, the measured intensity can be expressed by:

$$s'_0(\delta) = \frac{1}{2}(s_0 + s_1\cos(\delta) - s_3\sin(\delta)) \tag{6}$$

Four polarization states were measured with LCVR retardance values of $\delta = \{\pi/2, \pi, 3\pi/2, 2\pi\}$, corresponding to quarter-wave ($\lambda/4$), half-wave ($\lambda/2$), three-quarter-wave ($3\lambda/4$), and full-wave ($\lambda$) retardance, respectively. Note that for a vertically polarized laser source, retardances $\delta = \{\pi, 2\pi\}$ correspond to orthogonal horizontal and vertical linear polarizations (HLP and VLP), respectively. Analogously, $\delta = \{\pi/2, 3\pi/2\}$ correspond to orthogonal left and right circular polarizations (LCP and RCP), respectively.

The measurements result in four instances of (6), each comprised of $s_0$, $s_1$ and $s_3$ SHG Stokes vector components, therefore, the following measurement matrix can be defined:

$$\begin{pmatrix} s'_0(\pi/2) \\ s'_0(\pi) \\ s'_0(3\pi/2) \\ s'_0(2\pi) \end{pmatrix} = \frac{1}{2}\begin{pmatrix} 1 & 0 & -1 \\ 1 & -1 & 0 \\ 1 & 0 & 1 \\ 1 & 1 & 0 \end{pmatrix}\begin{pmatrix} s_0 \\ s_1 \\ s_3 \end{pmatrix} \tag{7}$$

To take advantage of all measured data, four subsystems of three equations were solved to compute $s_0$, $s_1$ and $s_3$ and the results were averaged to improve the robustness and accuracy. The remaining

$s_2$ component of $s'$ vector can be measured by probing elliptical polarization states with one LCVR or by introducing an additional LCVR in the PSA, however, computation of this component is omitted in this article.

The above analysis is repeated for each of the 4 incident laser polarization states, as prepared by the PSG using an infrared LCVR oriented at 45° with respect to the laser beam polarization. The PSG polarization states included the same retardances used in the PSA, corresponding to horizontal and vertical linear polarizations ($\delta = \{\pi, 2\pi\}$), and left and right circular polarizations ($\delta = \{\pi/2, 3\pi/2\}$), respectively. The resulting combination of 16 incident and outgoing polarization states provides a set of 12 Stokes vector elements, which can be expressed as:

$$s_{rDSMP} = \{s_0^{HLP}, s_1^{HLP}, s_3^{HLP}, s_0^{VLP}, s_1^{VLP}, s_3^{VLP}, s_0^{RCP}, s_1^{RCP}, s_3^{RCP}, s_0^{LCP}, s_1^{LCP}, s_3^{LCP}\} \quad (8)$$

, where the superscripts denote the PSG polarization state (incident polarization), and the subscripts indicate the component of the resulting SHG signal at the sample. These elements are then used to compute 5 polarimetric parameters: SHG intensity with circularly-polarized light, R-Ratio, degree of circular polarization (DCP), SHG circular dichroism (SHG-CD), and SHG linear dichroism (SHG-LD). The equations for the polarimetric parameters in terms of the Stokes vector components correspond to equations 2-6 in the "polarimetric parameter calculations" section of the Methods.

### Supplementary Note: Optimal data discretization for texture analysis

Texture analysis of the calculated polarimetric parameter images provides additional insight on the tissue ultrastructure in the form of a scalar score when computed over an image [4, 5, 6, 7]. A simple implementation of texture analysis over the whole image area could be performed; however, given the large field of view provided by widefield P-SHG microscopy, computed texture parameters would disregard the small-scale structural variations of the ECM. In order to highlight local texture variations of the tissue, polarimetric parameter images are subdivided, and texture analysis is performed on smaller sub-images. Investigations on the optimal number of sub-images is shown in Supplementary Note 3. In addition, mean and mean absolute deviation (MAD) of the polarimetric parameter values are calculated for each sub-image. Therefore, the resulting dataset includes the number of pixels corresponding to the SHG signal (pixel density or PD), as well as, statistics (mean and MAD) and texture (contrast, correlation, entropy, ASM, and IDM) of the sub-images of each widefield polarimetric parameter image, that are then used in classification and machine learning-assisted diagnostics.

There are 3 requirements for computation of the texture parameters: 1) level of discretization of the continuous polarimetric data (the so-called gray levels, $Ng$) [7], 2) The upper ($UB$), and 3) the lower bounds ($LB$) of the range of the input polarimetric parameter data.

In order to compute $Ng$ (requirement 1), each sub-image was discretized using the Freedman-Diaconis (FD) rule [8], which is commonly used to calculate the optimal bin width of histograms, $w_{FD} = 2n^{(-1/3)}IQR(x)$, where $IQR(x)$ is the interquartile range of the data ($x$), and $n$ is the number of points in the dataset. Using FD rule, $Ng$ can be expressed as:

$$Ng = \frac{\max(x) - \min(x)}{w_{FD}} \tag{9}$$

, where $\max(x)$ and $\min(x)$ denote the minimum and maximum of the polarimetric parameter distributions. The above procedure is then repeated for all present sub-images, resulting in a distribution of $Ng$ values. To ensure sufficient discretization of polarimetric parameters, the maximum of the $Ng$ distribution is considered, however, to avoid the effects of extreme outliers, the 99th percentile of the $Ng$ distribution was instead selected for texture analysis.

The $LB$ and $UB$ (requirements 2 and 3) were selected as the 1st and 99th percentile of each sub-image pixel values, respectively, to avoid extreme outliers. Similar to $Ng$, these computations are performed for all sub-images, resulting in $LB$ and $UB$ distributions. Furthermore, to correctly represent the true lower and upper bounds of the polarimetric parameters, the 1st and 99th percentiles of the $LB$ and $UB$ distributions were selected for texture analyses, respectively. It should be mentioned that $LB$ and $UB$ do not directly correspond to $\min(x)$ and $\max(x)$, respectively, since the latter denotes the absolute minimum and maximum of the polarimetric parameters.

It is important to examine the effects of image subdivision on tissue classification performance. Hence, the above calculations were performed at different subdivision levels, to determine optimal $Ng$, $LB$, and $UB$ for texture analysis performed at each subdivision level. It is evident that $Ng$ is different for each polarimetric parameter at lower subdivision levels (fewer sub-images per image)

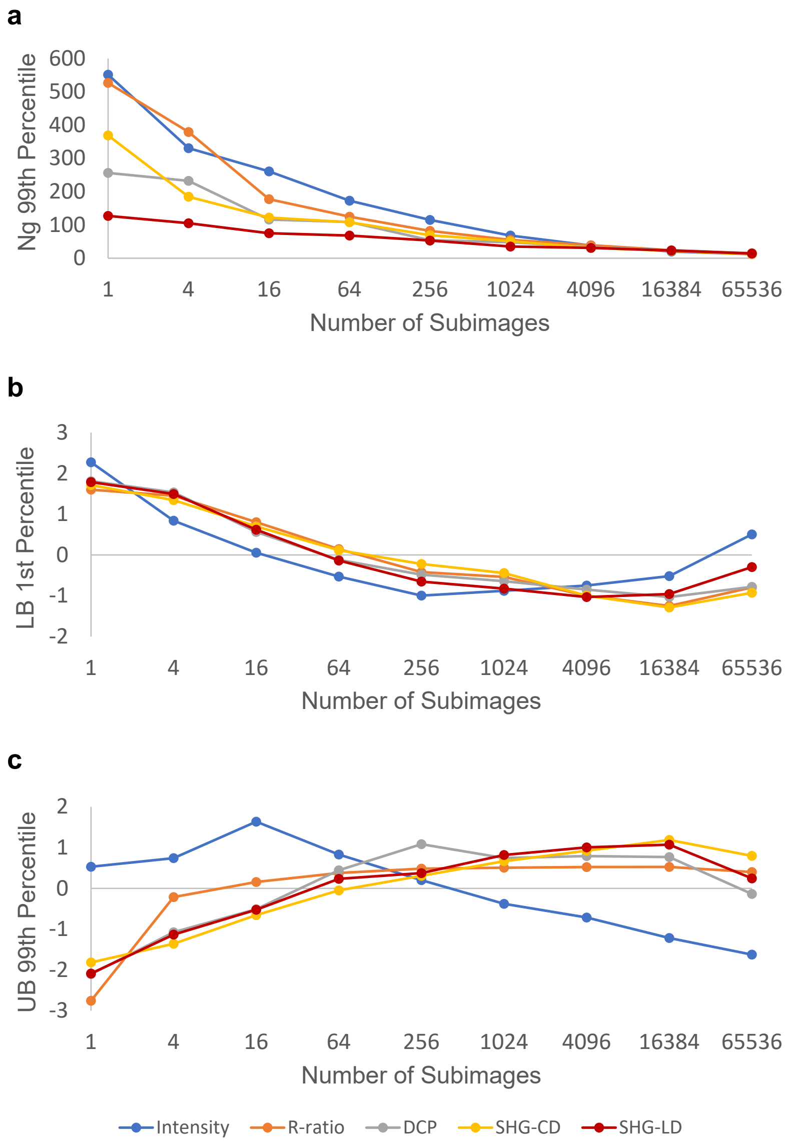


Figure 1: **Optimal discretization of continuous data for texture analysis**. a, Optimal number of gray levels ($Ng$) computed using Freedman-Diaconis rule for each subdivision level. 99th percentile of the $Ng$ distribution was used to represent maximum $Ng$. b-c, Lower ($LB$) and upper bounds ($UB$) of data, represented by the 99th and 1st percentiles of corresponding distributions, respectively. Curves are standardized for better visual comparisons.

and decreases with the number of sub-images. $Ng$ eventually converges to a value between 12 and 15 at the largest subdivision level due to limited pixel value variations over the small sizes of each sub-image (Supplementary Fig. 1a).

Standardized $UB$ and $LB$ curves are shown in Supplementary Fig. 1b-c for better comparison of the trends. The standardization involved subtraction of mean from each polarimetric parameters' $LB$ and $UB$, followed by division by their corresponding standard deviations. Overall, it is evident that $LB$ decreases, while $UB$ increases with the subdivision level. These trends are expected since most of the data are located near the center of the distribution. The SHG intensity exhibits highly right-skewed log-normal distributions, so that increasing the number of sub-images shortens the tail and decreases UB. The following section shows how optical number of sub-images can be found, using the computed $Ng$, $LB$, and $UB$.

#### Supplementary Note: Optimal number of sub-images

Once the normal and tumor training dataset are computed, the subdivision level was optimized to maximize classification performance. A series of classification techniques, including linear and quadratic discriminant analyses, as well as linear and quadratic support vector machines were used to differentiate between normal and tumor groups [9]. However, the highest degree of accuracy was achieved by a logistic regression model, which also provided easy-to-interpret classification probabilities, further enabling computation of an important threshold-independent performance metric, Brier score [10]. As such, a binary logistic regression classifier was trained using 1000x repeated 5-fold cross validation at different subdivision levels. The classifier performance was measured using threshold-dependent metrics (threshold at 50% posterior probability) such as accuracy, true negative and positive rates, and F1-score, as well as threshold-independent metrics, including the area under the receiver operating characteristic curve (AUROC) and Brier score [10, 11]. The mean and standard deviation of the repeated performance metric measurements were computed to assess the capability and stability of the classifier, as shown in Supplementary Fig. 2a-b.

It is evident that classification without subdivisions, corresponding to 1 sub-image per image, resulted in poor performance and low stability (highest standard deviation). The classification performance and stability were enhanced with increasing subdivision level. However, despite greatest stability at very high levels of subdivision, classification computation time was significantly lengthened (Supplementary Fig. 2d). Worse yet, a large difference between normal and tumor data size became apparent in high subdivision levels, which resulted in significant decrease of the true positive rate and the F1-score (Supplementary Fig. 2c). At high subdivision levels, the number of sub-images that did not possess SHG signal (thus discarded) increased disproportionately for tumor tissue due to sparseness of collagen fibers, resulting in the observed group data size disparity. As seen in Supplementary Fig. 2a, the true negative rate, corresponding to highly collagenous normal tissue, was unaffected by higher subdivision level and artificially increased the classification accuracy.

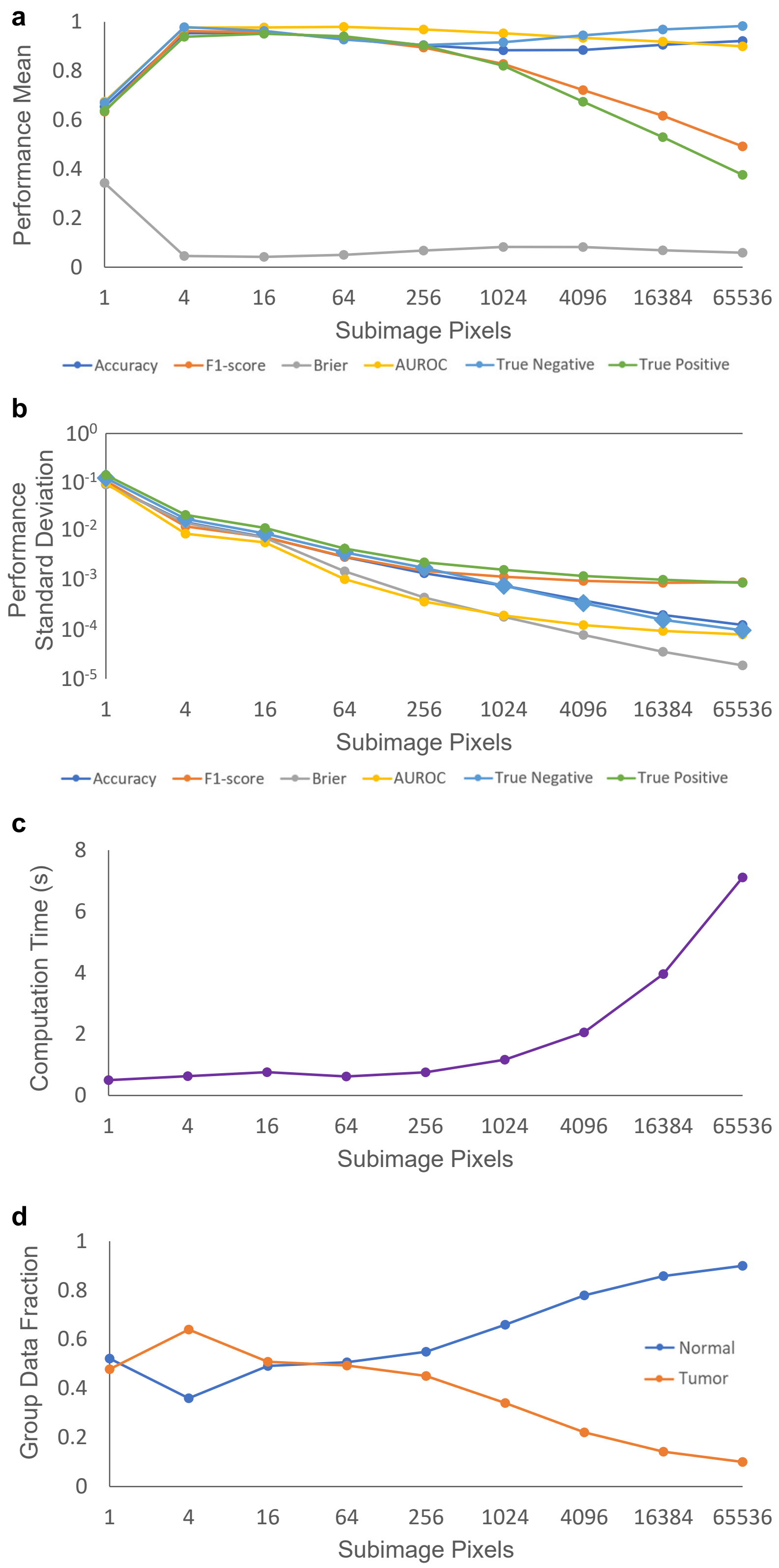


Figure 2: **Subdivision level optimization**. a, The mean of threshold-dependent metrics (accuracy, true positive and negative rates, and F1-score) and threshold-independent metrics (area under the receiver operating characteristic (AUROC) and Brier score) are calculated at various subdivision levels. b, Standard deviation of the performance metrics is computed to highlight classification stability at various subdivisions. It is evident stability increases rapidly with subdivision levels. c, Classification computation time increases quickly with subdivision levels beyond 256. d, Group data size disparity results from disproportionate omission of tumor sub-images with sparse collagen content, thus lacking SHG signal. This disparity has a direct effect on the true positive rate and the F1-score, as shown in a.

We determined the range of 4 to 64 sub-images per image was an acceptable subdivision level, possessing lowest Brier score and largest AUROC, and $> 90\%$ F1-score and true positive rate. Computation times are the lowest in this range, and the group size disparity is minimal, since normal and tumor groups each possess close to 50% of the training dataset (with the exception of 4 sub-images per image, in which case there are more tumor data points than normal which contributes to maintaining a high true positive rate). It is important to note that the classifier stability increases quickly to acceptable levels with the number of sub-images per image, reaching $< 5\%$ at 64 sub-images per image. For reasons stated above, 64 sub-images per image is chosen as the optimal subdivision level for classification of normal and tumor breast tissue in the manuscript.
